# DeepGO-SE: Protein function prediction as Approximate Semantic Entailment

**DOI:** 10.1101/2023.09.26.559473

**Authors:** Maxat Kulmanov, Francisco J. Guzmán-Vega, Paula Duek Roggli, Lydie Lane, Stefan T. Arold, Robert Hoehndorf

**Affiliations:** Computer, Electrical and Mathematical Sciences & Engineering Division, King Abdullah University of Science and Technology (KAUST), 4700 King Abdullah University of Science and Technology, Thuwal, 23955-6900, Makkah, Saudi Arabia; Bioscience Program, Biological and Environmental Science and Engineering Division, King Abdullah University of Science and Technology (KAUST), 4700 King Abdullah University of Science and Technology, Thuwal, 23955-6900, Makkah, Saudi Arabia; Computational Bioscience Research Center, King Abdullah University of Science and Technology (KAUST), 4700 King Abdullah University of Science and Technology, Thuwal, 23955-6900, Makkah, Saudi Arabia; CALIPHO group, SIB Swiss Institute of Bioinformatics, CMU, 1 rue Michel Servet, Geneva 4, 1211, Switzerland; Department of Microbiology and Molecular Medicine, Faculty of Medicine, University of Geneva, CMU, 1 rue Michel Servet, Geneva 4, 1211, Switzerland

**Keywords:** Protein Function, Gene Ontology, Ontology Embedding, Machine Learning

## Abstract

The Gene Ontology (GO) is one of the most successful ontologies in the biological domain. GO is a formal theory with over 100,000 axioms that describe the molecular functions, biological processes, and cellular locations of proteins in three sub-ontologies. Many methods have been developed to automatically predict protein functions. However, only few of them use the background knowledge provided in the axioms of GO for knowledge-enhanced machine learning, or adjust and evaluate the model for the differences between the sub-ontologies.

We have developed DeepGO-SE, a novel method which predicts GO functions from protein sequences using a pretrained large language model combined with a neuro-symbolic model that exploits GO axioms and performs protein function prediction as a form of approximate semantic entailment. We specifically evaluate DeepGO-SE on proteins that have no significant similarity with training proteins and demonstrate that DeepGO-SE can improve function prediction for those proteins.

## 1 Introduction

Protein function prediction is one of the key challenges in modern biology and bioinformatics as it enables better understanding of the roles and interactions of proteins within living systems. Accurate functional descriptions of proteins are necessary for tasks such as identification of drug targets, understanding disease mechanisms, and improving biotechnological applications in industry. While predicting protein structures has become increasingly accurate in recent years [1], predicting protein function remains challenging due to the small number of known functions combined with their complexity and interactions.

Functions of proteins are described using the Gene Ontology (GO) [2] which is one of the most successful ontologies in biology. GO includes three sub-ontologies for describing molecular functions (MFO) of a single protein, biological processes (BPO) to which proteins can contribute, and cellular components (CCO) where proteins are active. Researchers identify protein functions based on experiments and generate scientific reports which are then taken by database curators and added to knowledge bases. These annotations are generally propagated to homolog proteins. As a result, the UniProtKB/Swiss-Prot database [3] contains manually curated GO annotations for thousands of organisms and more than 550,000 proteins.

Recent protein function prediction methods rely on different sources of information such as sequence, interactions, protein tertiary structure, literature, coexpression, phylogenetic analysis, or the information provided in GO [4–20]. The methods may use sequence domain annotations [5, 6, 8, 11, 21], directly apply deep convolutional neural networks (CNN) [13] or language models such as LSTMs [9] and transformers [14], or use pretrained protein language models [10, 15] to represent amino acid sequences. Models may also incorporate protein–protein interactions through knowledge graph embeddings [12, 16], approaches using *k*-nearest neighbors [21], and graph convolutional neural networks [6]. Also, natural language models applied to scientific literature have been successful in automated function prediction [8].

One of the major limitations of many function prediction methods is their reliance on sequence similarity to predict functions. While this approach has been effective when applied to proteins that have similar proteins with well-characterized functions, it can be less reliable for proteins with little or no sequence similarity to known functional domains. Molecular functions arise largely from structure, and proteins with similar structures might have different sequence [22]. Importantly, proteins with similar sequences can have a different set of functions depending on their active sites and the organisms in which they are a part. Consequently, methods that use the same sources of information for all three sub-ontologies of GO are limited; while functions from the MFO sub-ontology can be predicted by a protein sequence or structure, functions from BPO and, to a lesser degree, CCO, inherently rely on multiple proteins being present and interacting in particular ways; therefore, predicting BPO and CCO annotations requires different sources of information than predicting MFO annotations. In general, predicting whether a protein participates in a biological process requires knowledge of an organisms proteome, or at least its annotated genome so that proteins can be predicted; as a result, two proteins may have 100% sequence identity but participate in different processes, depending on the presence or absence of other proteins within the organism’s proteome. Protein–protein interaction networks can encode the proteome as well as limit the search space for potential interactions between proteins that give rise to biological processes. Ontologies are another source of information rarely exploited for predicting protein functions. Ontologies are not simply collections of classes; rather, ontologies are formal theories that specify some aspects of the intended meaning of a class using a logic-based language [23]. The background knowledge that is contained in the axioms of GO can be used by some machine learning models to improve predictions through knowledge-enhanced machine learning [11, 12, 14, 15]. By incorporating the formal axioms into machine learning models, it becomes possible to leverage prior knowledge during the learning or prediction process, and to put constraints on the parameter search space that can improve the accuracy and efficiency of the learning process, and, ultimately, make better predictions [24, 25]. While there are different approaches of how formal background knowledge can be incorporated in machine learning methods, *approximate entailment* aims to explicitly and provably perform “semantic entailment” as optimization objective, and therefore reproduce many of the formal properties of deductive systems [26]. Only few function prediction methods utilize the formal axioms that are in GO. Hierarchical classification methods for predicting protein functions such as GoStruct2 [27], DeepGO [12] and DeePred [28] use subsumption axioms to extract hierarchical relations between classes but ignore the rest of the axioms stored in GO that can be used to reduce the search space and improve predictions.

We have developed DeepGO-SE, a novel protein function prediction method which predicts functions from protein sequences using a pretrained large protein language model combined with a neuro-symbolic model that performs function prediction as approximate semantic entailment. We use the ESM2 protein language model [29] to generate representations of single proteins. Similarly to DeepGOZero [11], we project the ESM2 embeddings into an embedding space (ELEmbeddings) that is generated from the axioms in the GO [30]. ELEmbeddings encode ontology axioms based on geometric shapes and geometric relations, and corresponds to a Σ algebra, or “world model”, in which we can determine whether statements are true or false. In contrast to DeepGOZero, we use these world models to perform “semantic entailment”: statement *φ* is entailed by theory *T* (*T* |= *φ*) if and only if *φ* is true in every world model in which all statements in *T* are true [31]. While there are, in general, infinitely many such world models for a theory *T* or a statement *φ*, we learn multiple, but finitely many, such models and generate predictions of functions as “approximate” semantic entailment where we test for truth in each of the generated world models. Using this form of approximate semantic entailment, we show that the axioms in the extended version of GO enhance the predictions of molecular functions.

Furthermore, we improve predictions for complex biological processes and cellular components by incorporating information about an organisms proteome and interactome in the form of protein–protein interaction networks. We show that, unlike molecular functions, predictions of annotations to biological processes and cellular components can significantly benefit from protein–protein interactions. For biological processes, we found that integrating predicted molecular functions and interactions considerably improves the performance of the predictions; this finding indicates that prediction of biological process annotations does not require knowledge of specific proteins but only their molecular functions, thereby substantially expanding the generality of our method.

We train and evaluate our model on a dataset with experimental annotations which is split based on sequence similarity to make sure that the evaluations are reported using a test set that does not share similar protein with the training set. We find that methods which rely on sequence similarity perform poorly in this setting, whereas DeepGO-SE significantly improves the prediction performance for all sub-ontologies of GO.

Overall, the contributions of our work are as follows:

- We developed a novel method for knowledge-enhanced machine learning as approximate semantic entailment over multiple generated world models.
- We developed a novel method for predicting protein functions which improves prediction performance of sub-ontologies of GO by using knowledge-enhanced learning and a combination of different sources of information.
- We improve function prediction performance for novel proteins by using sequence features generated by a pretrained protein language model ESM2.

## 2. Results

### 2.1 DeepGO Semantic Entailment

In the DeepGO-SE model, we use ESM2 [29] to represent protein sequence and project them into multiple geometric interpretations (i.e., models) of GO that have been generated with ELEmbeddings [30]; we then test the degree of truth of statements assigning a function to a protein in each interpretation of GO, and aggregate over all interpretations. The EMS2 embeddings of proteins are used as input to an MLP model that projects the embedding into ELEmbeddings space by matching the dimensionality of the EMS2 embedding with the dimension of the ELEmbedding space:

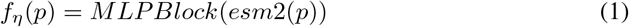

Within a model, given a protein *p* and GO class *c*, we test for the truth of the statement *p* ⊑ ∃hasFunction.*c* using the following formula:

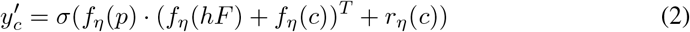

where *f*_*η*_(*p*) is the projection function, *f*_*η*_(*hF*) is the embedding of the *hasFunction* relation, *f*_*η*_(*c*) is the center embedding of an *n*-ball representing class *c, r*_*η*_(*c*) is the radius of the *n*-ball representing class *c*, and *Σ* is a sigmoid activation function.

The statement is approximately entailed if it is true in all interpretations generated by DeepGO-SE. We independently generate several ELEmbedding and projection functions *f*_*η*_(*p*), and combine the truth values for the tested axiom 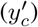 in each of the interpretations to obtain the final prediction scores (the degree of entailment). Given *N* interpretations, we combine truth values by computing the minimum, maximum, or arithmetic mean of the truth values in all *N* interpretations. Figure 1 provides an overview of the prediction model of DeepGO-SE.

**Fig. 1:**
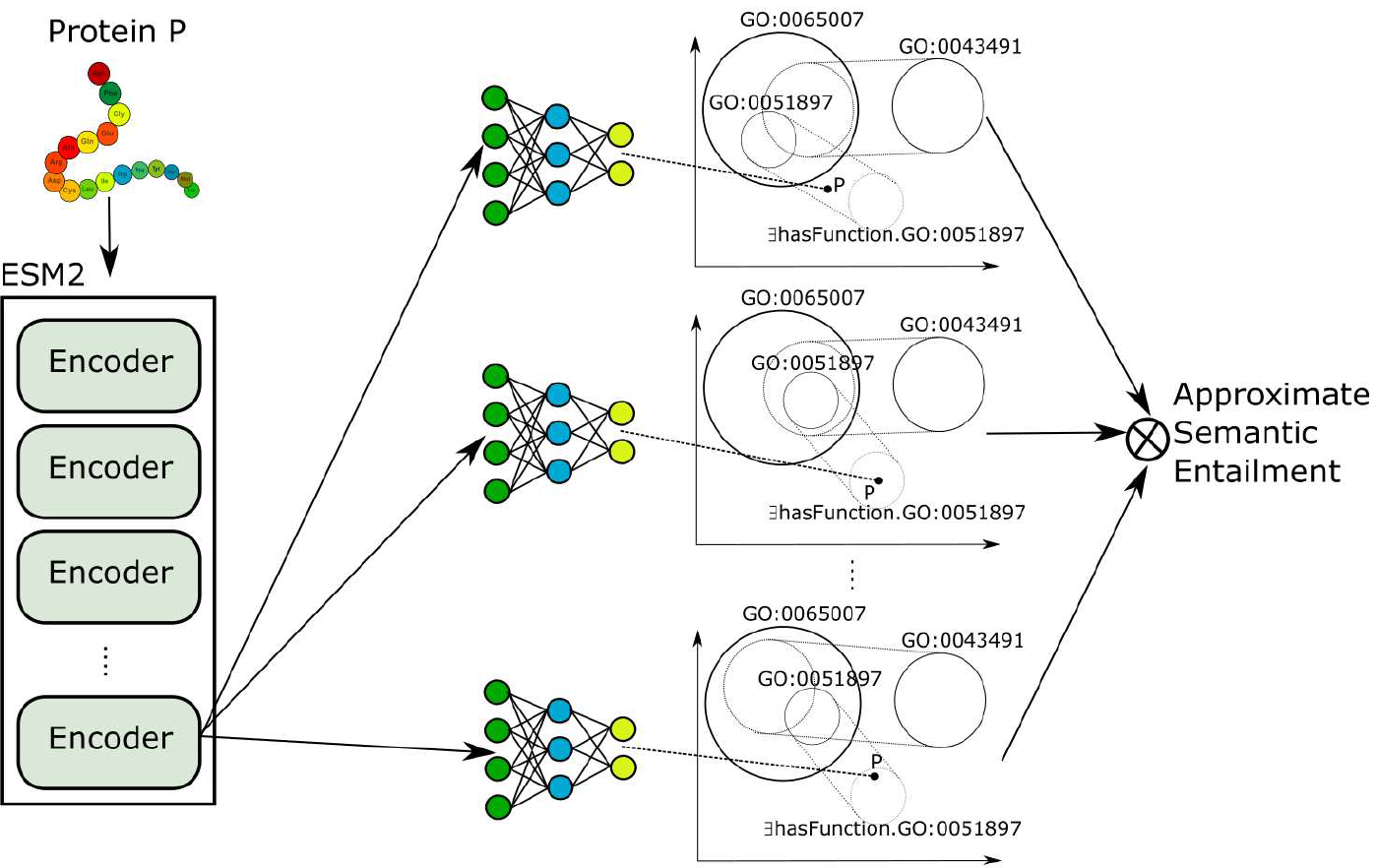
The figure provides a high-level overview of the DeepGO-SE model. On the left, a protein *P* is embedded in a vector space using ESM2 model whereas the right side shows multiple models with an MLP that embeds the protein in the same space as the GO axioms. Furthermore, predictions from multiple models are combined to perform approximate semantic entailment.

### 2.2 Evaluation of DeepGO-SE against the UniProtKB/Swiss-Prot dataset

We evaluate and compare our method with the baseline methods using the UniProtKB/Swiss-Prot dataset split by sequence similarity. We use protein-centric evaluation measures such as *F* max, *S*min, and AUPR, and class-centric AUC standardized by the CAFA challenge [32]. We provide detailed information about evaluation measures in the Supplementary materials.

We train and evaluate three sub-ontologies of GO separetely because they have different characteristics in terms of number of classes and their relations, number of proteins and sources of information they can benefit from. We compare with five baseline methods: Naïve, MLP, DeepGOCNN, DeepGOZero and DeepGraphGO. None of these methods relies on sequence similarity and, except for the naïve predictor, all assign functions based on sequence features that are learned directly or using features derived from tools such as InterProScan [33].

In all evaluations, the DeepGO-SE model significantly outperformed all the baseline methods in terms of *F* max, AUPR and AUC. In MFO, DeepGO-SE achieved an *F* max of 0.554 which is 7% larger than the result achieved by the MLP and DeepGOZero methods (Table 1). In predicting BPO annotations, the model achieves an *F* max of 0.432 which is around 8% higher than the best baseline method DeepGraphGO (Table 2), and in the CCO evaluation, DeepGO-SE model achieves *F* max of 0.721 (Table 3).

**Table 1:** Prediction results for Molecular Function on the UniProtKB/Swiss-Prot dataset.

| Method | $F_{\max}$ | $S_{\min}$ | AUPR | AUC |
| --- | --- | --- | --- | --- |
| Naive | 0.321 | 14.568 | 0.180 | 0.500 |
| MLP | 0.484 | 12.650 | 0.441 | 0.673 |
| DeepGOCNN | 0.404 | 13.741 | 0.365 | 0.749 |
| DeepGOZero | 0.483 | 12.722 | 0.444 | 0.749 |
| DeepGraphGO | 0.416 | 14.077 | 0.357 | 0.673 |
| DeepGO-SE | **0.554** | 11.681 | **0.552** | **0.874** |
| DeepGOGAT-SE | 0.525 | **11.137** | 0.523 | 0.861 |

This table shows protein-centric $F_{\max}$ , $S_{\min}$ , and AUPR, and the class-centric average AUC.

**Table 2:** Prediction results for Biological Process on the UniProtKB/Swiss-Prot dataset.

| Method | $F_{\max}$ | $S_{\min}$ | AUPR | AUC |
| --- | --- | --- | --- | --- |
| Naive | 0.294 | 43.934 | 0.195 | 0.500 |
| MLP | 0.350 | 42.059 | 0.303 | 0.668 |
| DeepGOCNN | 0.334 | 42.912 | 0.275 | 0.686 |
| DeepGOZero | 0.343 | 42.857 | 0.284 | 0.643 |
| DeepGraphGO | 0.354 | 42.100 | 0.303 | 0.736 |
| DeepGO-SE | 0.432 | 39.419 | 0.401 | 0.864 |
| DeepGOGAT-SE | 0.435 | 39.123 | 0.404 | **0.876** |
| DeepGOGATMF-SE | **0.448** | **37.299** | **0.428** | 0.831 |
| DeepGOGATMF-SE-Pred | 0.444 | 39.098 | 0.409 | 0.855 |

This table shows protein-centric $F_{\max}$ , $S_{\min}$ , and AUPR, and the class-centric average AUC.

**Table 3:** Prediction results for Cellular Component on the UniProtKB/Swiss-Prot dataset.

| Method | $F_{\max}$ | $S_{\min}$ | AUPR | AUC |
| --- | --- | --- | --- | --- |
| Naive | 0.620 | 11.879 | 0.490 | 0.500 |
| MLP | 0.623 | 11.584 | 0.622 | 0.636 |
| DeepGOCNN | 0.661 | 11.079 | 0.670 | 0.758 |
| DeepGOZero | 0.625 | 11.700 | 0.587 | 0.599 |
| DeepGraphGO | 0.667 | 10.020 | 0.666 | 0.814 |
| DeepGO-SE | 0.721 | 9.499 | 0.730 | 0.914 |
| DeepGOGAT-SE | **0.736** | **8.634** | 0.743 | **0.930** |
| DeepGOGATMF-SE | 0.668 | 9.809 | 0.679 | 0.884 |
| DeepGOGATMF-SE-Pred | 0.694 | 9.907 | **0.753** | 0.884 |

This table shows protein-centric $F_{\max}$ , $S_{\min}$ , and AUPR, and the class-centric average AUC.

In our basic DeepGO-SE model, protein embeddings are generated from protein sequence by ESM2; however, we can modify *f*_*η*_(*p*) to encode more information about a protein. We argue that biological process and cellular component annotations cannot be predicted from a protein sequence alone because even sequence-identical proteins can legitimately be involved in different processes dependent on the presence or absence of other proteins. Therefore, we use *f*_*η*_(*p*) to also encode information about a proteome and its interactions (protein– protein interactions, PPIs). To combine PPIs with individual features of proteins we use Graph Attention Networks (GAT) [34] and embed the protein *p* in the ELEmbeddings space using the formula

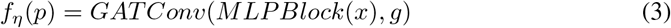

where *x* is an input feature vector for *p, g* is the PPI graph, *MLPBLock* is described in equation 6, *GATConv* is a GAT layer. We use this embedding function and alter the input vector *x* to perform three experiments. First, in DeepGOGAT-SE, we use the ESM2 embeddings as input for each protein. Second, in DeepGOGATMESE, *x* consists of the experimental annotations of a protein *p* to its molecular functions using a binary vector of size 6851. Third, in DeepGOGATMESE-Pred, we use the prediction scores from the DeepGO-SE model for molecular functions as input. We train and evaluate these three models to determine the effect of incorporating interactions.

Combining PPIs and ESM2 embeddings in the DeepGOGAT-SE model reduced the MFO prediction performance to an *F* max of 0.525, but slightly improved *S*min. Incorporating PPIs improves the performance in BPO predictions to *F* max of 0.435. The overall best performance in BPO is achieved when using experimental MFO annotations as features (DeepGOGATMF-SE), followed by MFO annotations predicted by DeepGO-SE (DeepGOGATMF-SE-Pred) (Table 2). For CCO, incorporating PPIs in the DeepGO-SE model increases *F* max from 0.721 to 0.736 (DeepGOGAT-SE) (Table 3).

Interestingly, including PPIs in our model did not improve MFO predictions (except for a slight improvement in *S*min), demonstrating that molecular functions can be predicted from single proteins whereas information about multiple proteins needs to be used to predict BPO and CCO annotations.

### 2.3 Evaluation of DeepGO-SE against the neXtProt manual prediction dataset

In order to further evaluate the performance of our method and baseline methods, we used a dataset of manually predicted protein functions from neXtProt. neXtProt assigns functions to uncharacterized proteins based on expert curation of available evidence. We found that, for molecular functions, the best *F* max of 0.386 is achieved by our the DeepGO-SE method and the second best *F* max 0.375 is achieved by DeepGOGAT-SE. Surprisingly, similar performance is achieved by the Naïve method which only uses the term frequency. However, when we evaluate based on AUPR and term centric AUC, we find that DeepGO-SE performs significantly better. The discrepancy can be explained by the small number of annotations. In this dataset, the median number of annotations is one, meaning that most proteins have only one specific GO function prediction (Table 4).

**Table 4:** Prediction results for Molecular Function on the neXtProt dataset.

| Method | $F_{\max}$ | $S_{\min}$ | AUPR | AUC |
| --- | --- | --- | --- | --- |
| Naive | 0.360 | 10.340 | 0.165 | 0.500 |
| MLP | 0.370 | 10.555 | 0.298 | 0.516 |
| DeepGOCNN | 0.348 | 10.641 | 0.270 | 0.599 |
| DeepGOZero | 0.337 | 10.662 | 0.261 | 0.573 |
| DeepGraphGO | 0.330 | 10.573 | 0.270 | 0.558 |
| DeepGO-SE | **0.386** | **10.093** | **0.324** | **0.744** |
| DeepGOGAT-SE | 0.375 | 10.254 | 0.291 | 0.700 |

This table shows protein-centric $F_{\max}$ , $S_{\min}$ , and AUPR, and the class-centric average AUC.

For biological processes, our method DeepGOGAT-SE which combines PPIs into the model performs with the best *F* max of 0.350. DeepGO-SE achieves slightly lower *F* max of 0.349 and slightly better *S*min, however DeepGOGAT-SE is substantially better in terms of AUPR and AUC. The third best *F* max and the best AUC is achieved by our method which uses predicted molecular functions to predict biological processes. We were not able to evaluate the DeepGOGATMF-SE method because many of the proteins are missing manual molecular function (Table 5).

**Table 5:** Prediction results for Biological Process on the neXtProt dataset.

| Method | $F_{\max}$ | $S_{\min}$ | AUPR | AUC |
| --- | --- | --- | --- | --- |
| Naïve | 0.308 | 32.987 | 0.183 | 0.500 |
| MLP | 0.311 | 32.140 | 0.251 | 0.580 |
| DeepGOCNN | 0.286 | 32.152 | 0.235 | 0.571 |
| DeepGOZero | 0.329 | 31.999 | 0.263 | 0.553 |
| DeepGraphGO | 0.322 | 31.861 | 0.240 | 0.558 |
| DeepGO-SE | 0.349 | **30.170** | **0.312** | 0.683 |
| DeepGOGAT-SE | **0.350** | 30.218 | **0.312** | 0.666 |
| DeepGOGATMF-SE-Pred | 0.339 | 30.653 | 0.293 | **0.694** |

This table shows protein-centric $F_{\max}$ , $S_{\min}$ , and AUPR, and the class-centric average AUC.

### 2.4 Case study: Validation based on structural homologs

We further investigated some predictions of molecular functions for which DeepGO-SE and neXtProt were in agreement to test if we could find additional evidence for the predictions. Specifically, we investigated the Mab-21-like protein 4 (MAB21L4) protein which has a single MFO annotation *nucleotidyltransferase activity* (`GO:0016779`), which is both assigned by neXtProt and DeepGO-SE. DeepGO-SE predicts this annotation with a high score of 0.638. MAB21L4 was predicted by neXtProt to be a nucleotidyltransferase based on available information about the protein’s activity in epidermal keratinocytes [35]. As part of investigating the role of MAB21L4 in keratinocytes, distant homology detection was used to assign MAB21L4 to the nucleotidyltransferase (NTase) fold superfamily [36]. The active site is described by the motifs hG[GS], [DE]h[DE]h and h[DE]h (h indicates a hydrophobic amino acid), where the three conserved aspartate/glutamate are involved in coordination of divalent ions and activation of acceptor hydroxyl group of the substrate, and the hG[GS] pattern is involved in holding the substrates within the active site [36]. Sequence alignment combined with structural data provided by AlphaFold suggests that the [DE]h[DE]h motif is conserved in MAB21L4 (Asp80-Met81-Glu-82-Val83) while the h[DE]h motif that aligns with other family members may not be conserved as it is replaced by an histidine (Phe199-His200-Val201). An alternative h[DE]h motif is present in Val-236, Asp-237, Leu-238 with the two first residues in a loop and the third one at the beginning of a short *β*-strand. The hG[GS] motif is less conserved among the nucleotidyltransferase superfamily members and seems not conserved among the members of the Mab-21 group but is present in Mab-21-like protein 1 (MAB21L1); sequence-based methods like InterProScan identify a *Mab-21-like, nucleotidyltransferase domain* (`IPR046903`) in MAB21L1. We used foldseek [37] to compare MAB21L1 and MAB21L4 structurally, and find that both are structurally very similar despite a low sequence similarity. Furthermore, MAB21L4 is structurally very similar to Cyclic GMP-AMP synthase (CGAS), which is well-characterized as having the *nucleotidyltransferase activity*.

Another noticeable example is the Family With Sequence Similarity 151 Member B (FAM151B) protein which was predicted to be a phosphoric diester hydrolase (`GO:0008081`) based on structural similarity to a protein from *Sicarius terrosus* by the neXtProt database. DeepGO-SE predicted the same function with a high score of 0.846. Foldseek search resulted in many sequence and structure homologs. Structure homologs with high sequence identity were not annotated, however, we found several well annotated structural homologs with low sequence identity. For example, the human protein Lysophospholipase D (GDPD3) has a high structural similarity to FAM151B and has been annotated with *phosphoric diester hydrolase activity* (`GO:0008081`) based on experimental evidence (Supplementary Figure B1). In addition, DeepGO-SE predicts other functions such as *metal ion binding* (`GO:0046872`) which GDPD3 has been annotated with as well. These findings suggest that DeepGO-SE learned to predict functions, among others, based on structural information.

### 2.5 Ablation study

In order to evaluate the contribution of the individual components of our models, we performed an ablation study. First, for each of the models, we removed the ElEmbeddings axiom loss functions and only optimized function prediction loss to determine the effect of using background knowledge contained in the GO. In the DeepGO-SE model, removing axioms losses resulted in a performance drop in the MFO evaluation while the performance in the BPO and CCO evaluations was not affected. Second, we trained the models with only GO or only GO-PLUS axioms to further evaluate the effect of using more background knowledge for performing approximate semantic entailment. We found that the performance of the MFO model improves with GO-PLUS axioms compared to GO axioms whereas the performance of the BPO and CCO models slightly drop when using the additional axioms contained in GO-PLUS.

Using PPI information, in the DeepGOGAT-SE model, removing axioms and removing the semantic entailment module resulted in a slight performance increase in MFO evaluation but the performance dropped in the BPO and CCO evaluations. In models that use PPIs and molecular functions as protein features, performance is better for BPO and CCO when removing axioms and semantic entailment.

Overall, the ablation study shows that the ontology axioms and semantic entailment mostly contribute to MFO and CCO model performance whereas the performance of BPO model is not significantly affected. The PPIs with GAT noticably contribute to CCO and BPO model performance and BPO model achieves the best performance without axioms and semantic entailment. Supplementary Table B5 provides the results of the ablation study for all four evaluation measures.

## 3. Methods

### 3.1 UniProtKB/Swiss-Prot Dataset

We use a dataset that was generated from manually curated and reviewed dataset of proteins from the UniProtKB/Swiss-Prot Knowledgebase [3] version 2021 04 released on 29-Sep-2021. We filtered all proteins with experimental functional annotations with evidence codes EXP, IDA, IPI, IMP, IGI, IEP, TAS, IC, HTP, HDA, HMP, HGI, HEP. The dataset contains 77, 647 reviewed and manually annotated proteins. For this dataset we use Gene Ontology (GO) released on 2021-11-16. We train and evaluate models for each of the sub-ontologies of GO separately.

We mainly aim to predict functions of novel proteins that have a low sequence similarity to existing proteins in the dataset. Therefore, we decided to split our dataset based on any similarity hit with maximum e-value score of 0.001. We computed pairwise similarity using Diamond (v2.0.9) [38] and grouped the sequences that have some similarity and split these groups into training, validation and testing sets. Table 6 summarizes the datasets for each sub-ontology. We train and evaluate a separate model for each sub-ontology.

**Table 6:** Summary of the UniProtKB/Swiss-Prot dataset.

| Ontology | GO Terms | Proteins | Groups | Training | Validation | Testing |
| --- | --- | --- | --- | --- | --- | --- |
| MFO | 6,851 | 43,279 | 6,963 | 52,072 | 2,964 | 4,221 |
| BPO | 21,356 | 58,729 | 9,463 | 52,584 | 2,870 | 3,275 |
| CCO | 2,829 | 59,257 | 10,019 | 48,318 | 4,970 | 5,969 |

The table shows the number of GO terms, total number of proteins, number of groups of similar proteins, number of proteins in training, validation and testing sets for the UniProtKB/Swiss-Prot dataset.

### 3.2 neXtProt Dataset

In order to further evaluate the performance of our models we use a dataset of manually annotated predictions of uncharacterized human proteins from the neXtProt [39] database. neXtProt standardizes and integrates information on human proteins and provides users with an advanced search capability built around semantic technologies [39]. neXtProt contains free text summaries of the literature and standardized enzyme annotations from UniProtKB/Swiss-Prot, pathway annotations from KEGG [40] and Reactome [41], and GO MFO and BPO terms from a variety of resources, obtained either manually or by automatic procedures, and based on either experiments or computational analysis. Proteins lacking the above-mentioned annotations and those that are solely annotated with broad GO terms are considered as uncharacterized. They can be retrieved using the SPARQL [42] query `NXQ 00022` [43]. In the 2023-04-18 release of neXtProt, there are 1,521 such proteins. To stimulate the characterization of these poorly-studied proteins, neXtProt collects and reviews functional predictions from the literature and proposes their own function annotations based on a manual interpretation of different types of public data (phenotypes, expression, subcellular localization, protein and genetic interactions, phylogeny, structure, sequence and functional assays) [44]. These predictions are displayed in the function prediction pages as GO MFO or BPO terms, and the underlying evidence using the Evidence Code Ontology (ECO) [45].

Here we use the data retrieved from 113 publications together with different resources that were used to predict the functions of 239 uncharacterized human proteins. In total, the proteins collected 659 specific GO function annotations where 69 molecular functions were assigned to 53 proteins and 590 biological processes were assigned to 225 proteins. Roughly one third of the proteins (38%) are assigned to only one function that in most of the cases (85%) is a GO BPO term. Most of the functional predictions (78%) are based on one piece of evidence.

### 3.3 Protein Language Model ESM2

Protein language models are large transformer architectures trained on protein sequences. The Evolutionary Scale Model (ESM) [29, 46] has been trained on 250 million sequences and learned protein sequence representations that are predictive for biochemical and biological properties of proteins including their functions. The second version of ESM has been improved to learn better representations that are also predictive of tertiary structures of proteins. We use the pretrained model of ESM2 with 3 billion parameters (esm2 t36 3B UR50D) to generate representations of proteins in our dataset. For a protein, we compute the output of the last layer and take the mean of embeddings for each amino acid, resulting in an embedding of size of 2560 for each protein.

### 3.4 GO-PLUS

The standard version of GO does not include relations between GO classes and external ontologies such as ChEBI [47], Uberon [48], the Cell Ontology [49], or to structured vocabularies such as the NCBI Taxonomy [50]. These relations and cross-ontology axioms exist in an extended version called GO-PLUS [51]. For example, in GO-PLUS a class *Atrioventricular bundle cell differentiation* (`GO:0003167`) is defined as equivalent to *Cell differentiation* (`GO:0030154`) and *results in acquisition of features of* (`RO:0002315`) some *Atrioventricular bundle cell* (`CL:0010005`). We use the GO-PLUS ontology version released on 2021-11-16 which has over 260K axioms. Like GO, GO-PLUS uses the Web Ontology Language (OWL) 2 [52] to represent its axioms. The Description Logic fragment of OWL 2, OWL 2 DL, defines several profiles, i.e., restricted languages with specific computational properties. GO is formalized using the OWL EL profile [53]. However, GO-PLUS contains axioms that are not part of the OWL EL profile; therefore, it cannot directly be used with reasoning or machine learning methods that are based on OWL EL. We identify around 1, 500 axioms that do not fit in the OWL EL profile and filtered them out using the EL Vira tool [54].

### 3.5 Approximate Semantic Entailment

Suppose 𝒪 is an ontology composed of a set of class symbols C, relation symbols R, and individual symbols I, and that it is expressed in the Description Logic 𝒜ℒ𝒞 [55]. In this logic, each class symbol is considered a class description. If *C* and *D* are class descriptions and *R* is a relation symbol, then the expressions *C* ⊓ *D, C* ⊔ *D*, ¬*C*, ∀*R*.*C*, and ∃*R*.*C* are also considered as class descriptions.

In the 𝒜ℒ𝒞 Description Logic, axioms can be classified as TBox or ABox axioms. If *C* and *D* are class descriptions, *a* and *b* are individual symbols, and *r* is a relation symbol, a TBox axiom has the form *C* ⊑ *D*, while an ABox axiom has the form *C*(*a*) or *r*(*a, b*). A TBox is a set of TBox axioms, and an ABox is a set of ABox axioms. An interpretation ℐ = (Δ^*ℐ*^, ·^*ℐ*^) in 𝒜ℒ𝒞 comprises a non-empty domain Δ^*ℐ*^ and an interpretation function ·^*ℐ*^ that satisfies *C*^*ℐ*^ ⊆ Δ^*ℐ*^ for all *C* ∈ C, *R*^*ℐ*^ ⊆ Δ^*ℐ*^ × Δ^*ℐ*^ for all *R* ∈ R, and *a*^*ℐ*^ ∈ Δ^*ℐ*^ for all *a* ∈ I. The interpretation function is extended to concept descriptions as follows:

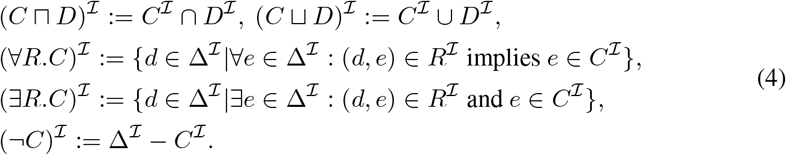

An interpretation ℐ is called a model of a TBox if, for all *C* ⊑ *D* in the TBox, *C*^*ℐ*^ ⊆ *D*^*I*^; and a model of an ABox if, for all *R*(*a, b*), (*a*^*ℐ*^, *b*^*ℐ*^) ∈ *R*^*I*^ and for all *C*(*a*), *a*^*ℐ*^ ∈ *C*^*ℐ*^.

A statement *φ* is semantically entailed by ontology 𝒪 (consisting of TBox and ABox), denoted *O* |= *φ*, if and only if every model of 𝒪 (i.e., an interpretation ℐ that is a model of both ABox and TBox of 𝒪) is also a model of *φ*(*Mod*(𝒪) ⊆ *Mod*(*φ*)). Semantic entailment requires access to all models of 𝒪 and *φ* which are usually infinite; approximate semantic entailment considers only a strict (usually finite) subset of *Mod*(𝒪) and tests whether *φ* is true in each of them [26, 56].

Here, we perform approximate semantic entailment by learning several models and determining whether a prediction (i.e., a statement that assigns a function to a protein) is true in all of them. For each sub-ontology of GO we train up to 10 models and combine the prediction scores using three different strategies. First, we take the maximum of the selected scores which means that if the predictions is made, it is true in all models. Second, we take an average of the scores. Here, the prediction is made if the prediction threshold is lower than average of all models. Lastly, we take the minimum of the scores where we make sure that the prediction is true in at least one of the selected models. We select the best parameters of the approximate semantic entailment based on our validation set and use the same on our test set. Supplementary Table B1 summarizes the results of semantic entailment on our validation set.

### 3.6 Protein–protein interaction (PPI) networks

Molecular functions of proteins mainly depend on their sequences and structures. However, biological processes result from interactions between multiple proteins. Therefore, to accurately predict biological processes, it is necessary to include multiple proteins and their interactions.

For our experiments, we use functional interactions between proteins provided by the STRING database (v11.0). We filter out all the interactions with confidence score less than 700. Our dataset uses UniProtKB identifiers and we map them to STRING database identifiers with mappings provided by UniProtKB. We generate the protein interaction graph using all the proteins in our dataset and use the DGL [57] library to process it and train graph neural networks.

### 3.7 Baseline methods

For our evaluations we selected methods that do not rely on predictions based on sequence similarity because our aim is to test the predictors on novel sequences. Therefore, we do not include methods as baselines that are primarily based on sequence similarity, such as predictions using BLAST or Diamond, or any other predictors that use their combinations.

3.7.1 Naive approach

Due to the imbalance in GO class annotations and propagation based on the true-path-rule, some classes have more annotations than others. Therefore, it is possible to obtain prediction results just by assigning the same GO classes to all proteins based on annotation frequencies. In order to test the performance obtained based on annotation frequencies, CAFA introduced a baseline approach called “naive” classifier [32]. Here, each query protein *p* is annotated with the GO classes with a prediction scores computed as:

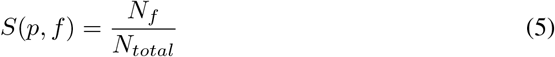

where *f* is a GO class, *N*_*f*_ is a number of training proteins annotated by GO class *f* and *N*_*total*_ is a total number of training proteins. We implement the same method.

3.7.2 MLP

The MLP baseline method predicts protein functions using a multi-layer perceptron (MLP) from a protein’s InterPro domain annotations obtained with InterProScan [33]. We represent a protein with a binary vector for all the InterPro domains and pass it to two layers of MLP blocks where the output of the second MLP block has residual connection to the first block. This representation is passed to the final classification layer with sigmoid activation function.

One MLP block performs the following operations:

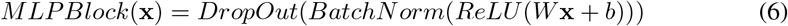

The input vector x of length 26, 406 represents InterPro domain annotations and is reduced to 1, 024 by the first MLPBLock:

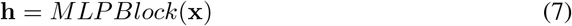

This representation is passed to the second MLPBlock with the input and output size of 1, 024 and added to itself using residual connection:

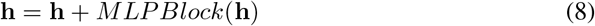

Finally, we pass this vector to a classification layer with a sigmoid activation function. The output size of this layer is the same as the number of classes in each sub-ontology:

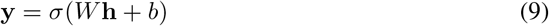

We train a different model for each sub-ontology in GO.

3.7.3 DeepGO-PLUS and DeepGOCNN

DeepGO-PLUS [13] predicts function annotations of proteins by combining DeepGOCNN, which predicts functions from the amino acid sequence of a protein using a 1-dimensional convolutional neural network (CNN), with the DiamondScore method. DeepGOCNN captures sequence motifs that are related to GO functions. Here, we only use CNN based predictions.

3.7.4 DeepGOZero

DeepGOZero [11] combines protein function prediction with a model-theoretic approach for embedding ontologies into a distributed space, ELEmbeddings [30]. ELEmbeddings represent classes as *n*-balls and relations as vectors to embed ontology semantics into a geometric model. It uses InterPro domain annotations represented as binary vector as input and applies two layers of MLPBlock as in our MLP baseline method to generate an embedding of size 1024 for a protein. It learns the embedding space for GO classes using ELEmbeddings loss functions and optimizes together with protein function prediction loss.

3.7.5 DeepGraphGO

The DeepGraphGO [6] method uses a neural network to combine sequence features (Inter-PRO domain annotations) with PPI networks by using graph convolutional neural networks. We have implemented DeepGraphGO based on the manuscript and provide the source code for our implementation. We trained and evaluated the model using our dataset.

## 4. Discussion

DeepGO-SE is a protein function prediction method that improves prediction performance for proteins by incorporating both protein sequence features generated by a pretrained protein language model, background knowledge from the GO, and interactions between proteins. Our results allow us to draw three main conclusions: knowledge-enhanced machine learning methods are now able to improve over methods that do not rely on background knowledge; GO function prediction is best formulated using a separate, hierarchical prediction approach; and function prediction models based on ESM2 can now generalize to largely unseen proteins. Although DeepGO-SE can predict biological processes and cellular components using only a protein sequence, the best performance is achieved when the sequence is combined with PPIs. However, many novel proteins do not have known interactions which limits the application of the combined model on them. Therefore, there is a need for methods which can accurately predict PPIs for novel proteins based on only available sequence. In future, we plan to incorporate sequence ans structure based PPI predictors into our model.

In addition, DeepGO-SE is able to perform zero-shot predictions, similar to DeepGOZero, and is faster to obtain predictions than other methods that rely on similarity search or multiple sequence alignments. This is due to the fact that DeepGO-SE relies only on ESM2 embeddings, which are faster to compute. Overall, the DeepGO-SE model represents a significant improvement over existing protein function prediction methods, providing a more accurate, comprehensive, and efficient approach.

## Acknowledgments

For computer time, this research used the resources of the Super-computing Laboratory at King Abdullah University of Science & Technology (KAUST) in Thuwal, Saudi Arabia.

## Availability of data and materials

All data underlying this work is freely available at https://github.com/bio-ontology-research-group/deepgo2.

## Code availability

The source code of this work is freely available at https://github.com/bio-ontology-research-group/deepgo2.

## Authors’ contributions

RH and MK conceived and designed the research. RH led the study and MK developed the prediction model and evaluated it. PD and LL contributed to functional analysis. FJGV and STA contributed to structural analysis. RH and MK developed the first draft of the manuscript. All authors contributed to writing and improving the manuscript and approved the submission.

## Appendix A Evaluation metrics

We use four different measures to evaluate the performance of our models. Three protein-centric measures *F*_max_, *S*_min_ and AUPR and one class-centric AUC.

*F*_max_ is a maximum protein-centric F-measure computed over all prediction thresholds. First, we compute average precision and recall using the following formulas:

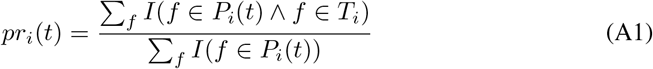

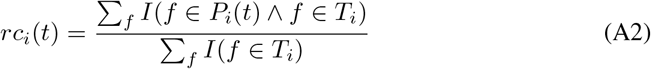

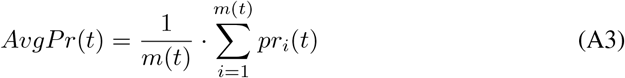

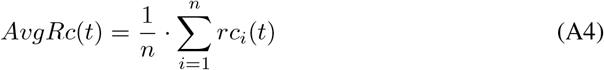

where *f* is a GO class, *T*_*i*_ is a set of true annotations, *P*_*i*_(*t*) is a set of predicted annotations for a protein *i* and threshold *t, m*(*t*) is a number of proteins for which we predict at least one class, *n* is a total number of proteins and *I* is an indicator function which returns 1 if the condition is true and 0 otherwise. Then, we compute the *F*_max_ for prediction thresholds *t* ∈ [0, 1] with a step size of 0.01. We count a class as a prediction if its prediction score is greater or equal than *t*:

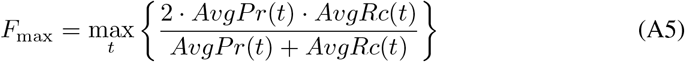

*S*_min_ computes the semantic distance between real and predicted annotations based on information content of the classes. The information content *IC*(*c*) is computed based on the annotation probability of the class *c*:

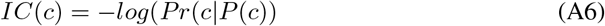

where *P* (*c*) is a set of parent classes of the class *c*. The *S*_min_ is computed using the following formulas:

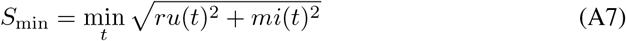

**Fig. B1:**
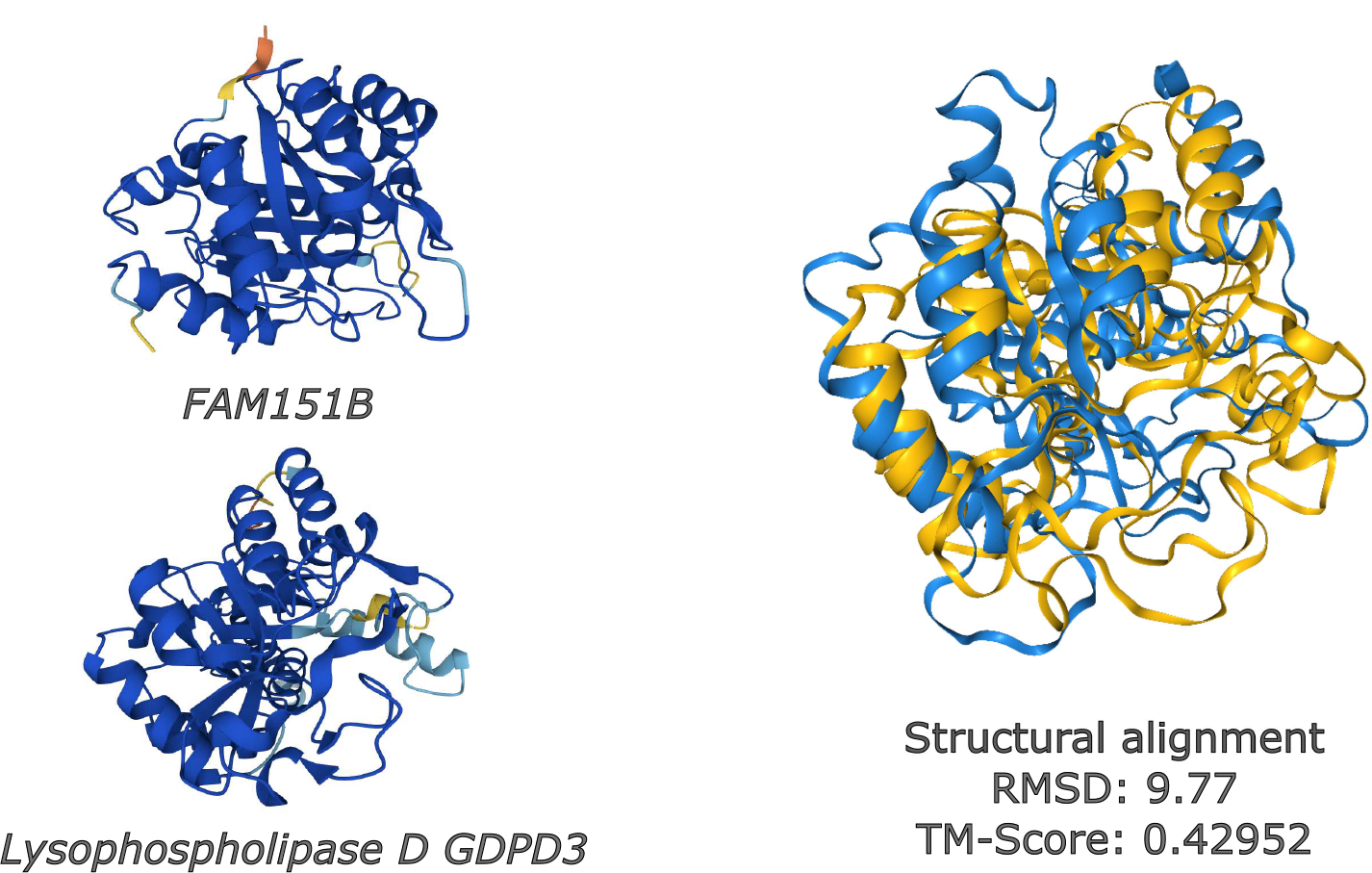
The figure provides structural comparison between human proteins *FAM151B* and *Lysophos-pholipase D GDPD3*. Foldseek search score is 1.0, RMSD is 9.77 and TM-Score is 0.42952.

where *ru*(*t*) is the average remaining uncertainty and *mi*(*t*) is average misinformation:

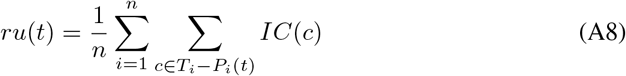

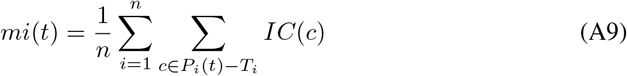

AUPR is the area under the average precision (*AvgPr*) and recall (*AvgRc*) curve.

AUC is a class-centric measure where compute AUC ROC per each class and take the average.

## Appendix B Supplementary figures

## Appendix C Supplementary tables

**Table C1:**
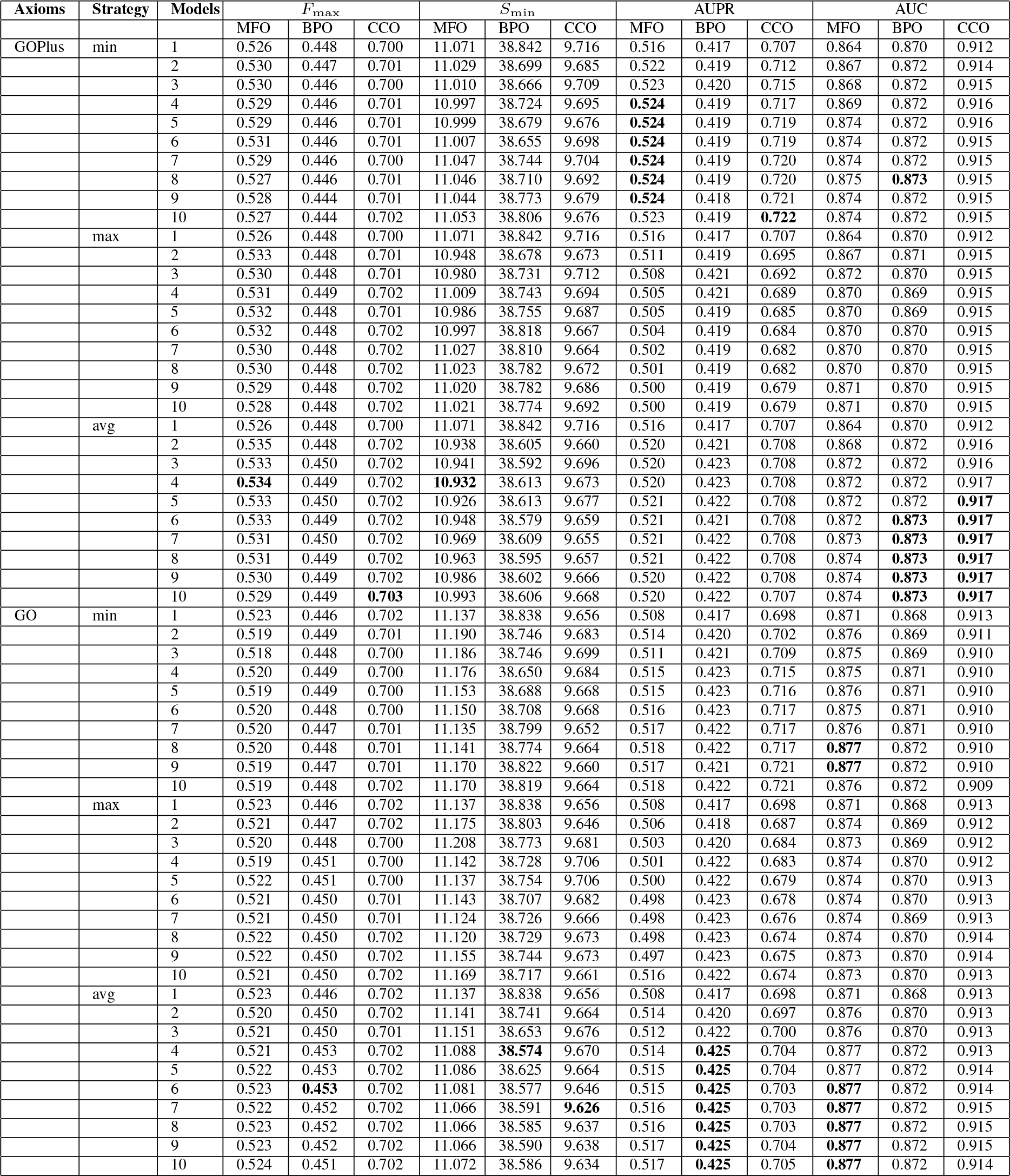
Performance of semantic entailment depending on axioms, combination strategy and number of models for Deep-GOSE model.

**Table C2:**
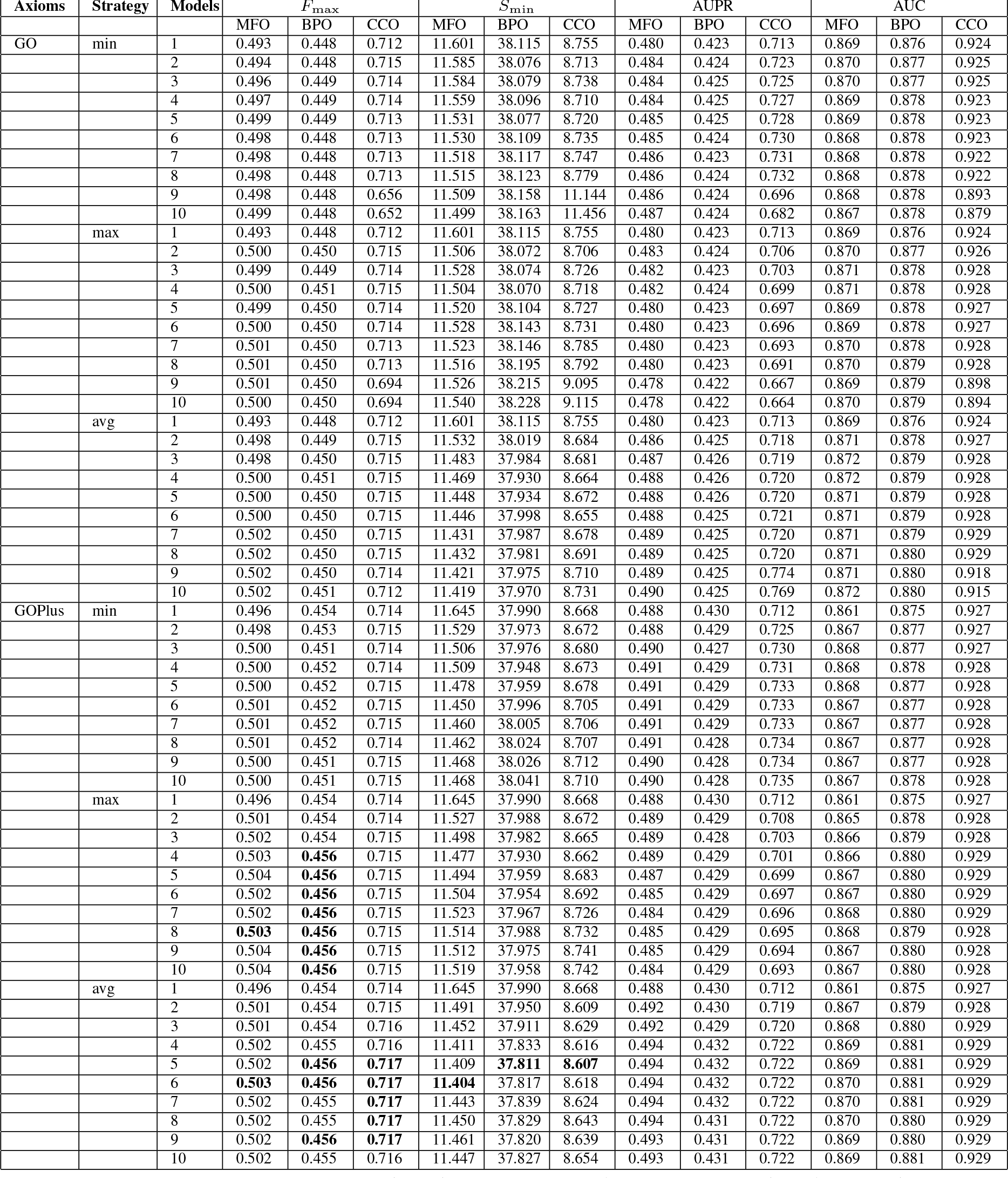
Performance of semantic entailment depending on axioms, combination strategy and number of models for Deep-GOGATSE model.

**Table C3:**
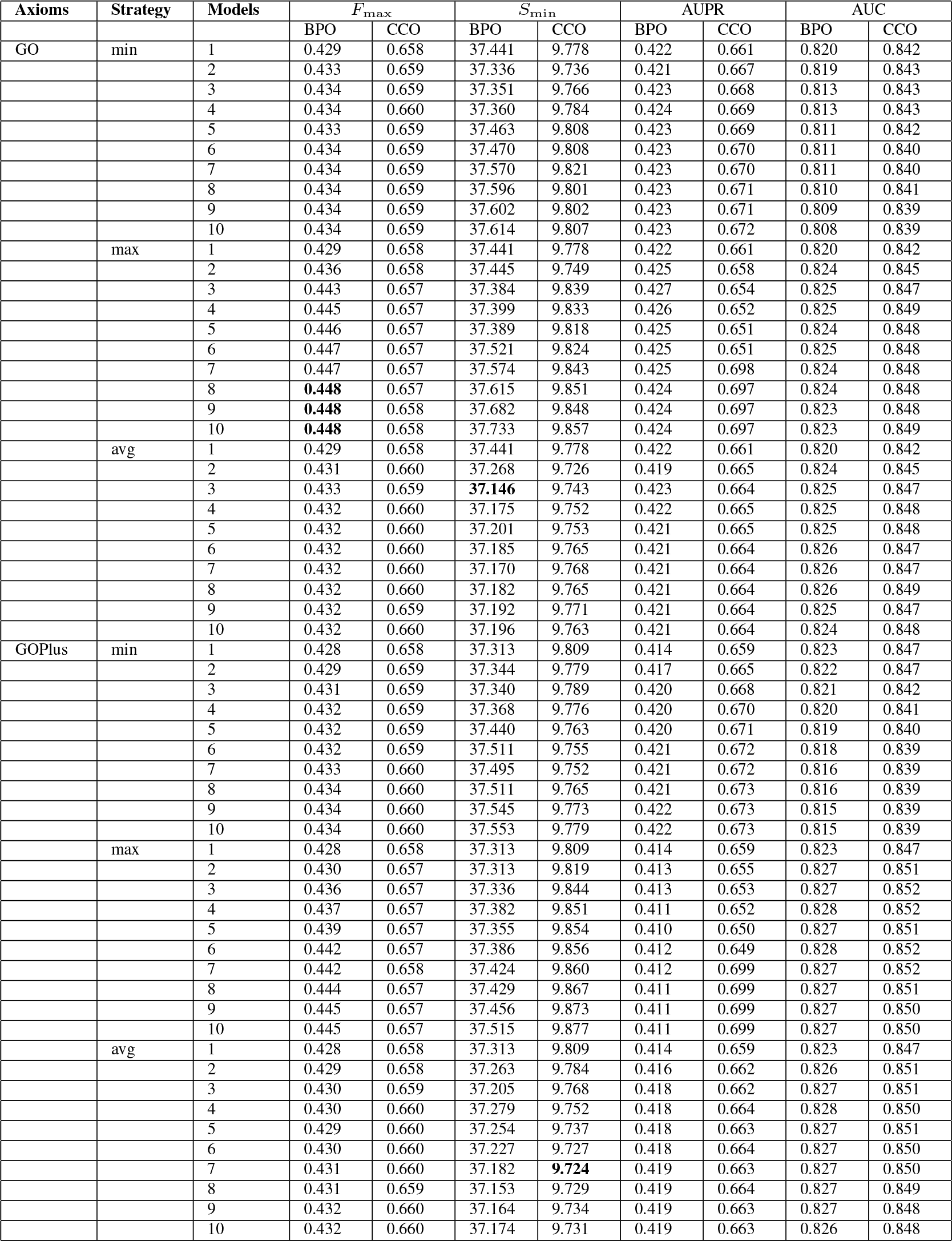
Performance of semantic entailment depending on axioms, combination strategy and number of models for DeepGOGATMFSE model.

**Table C4:**
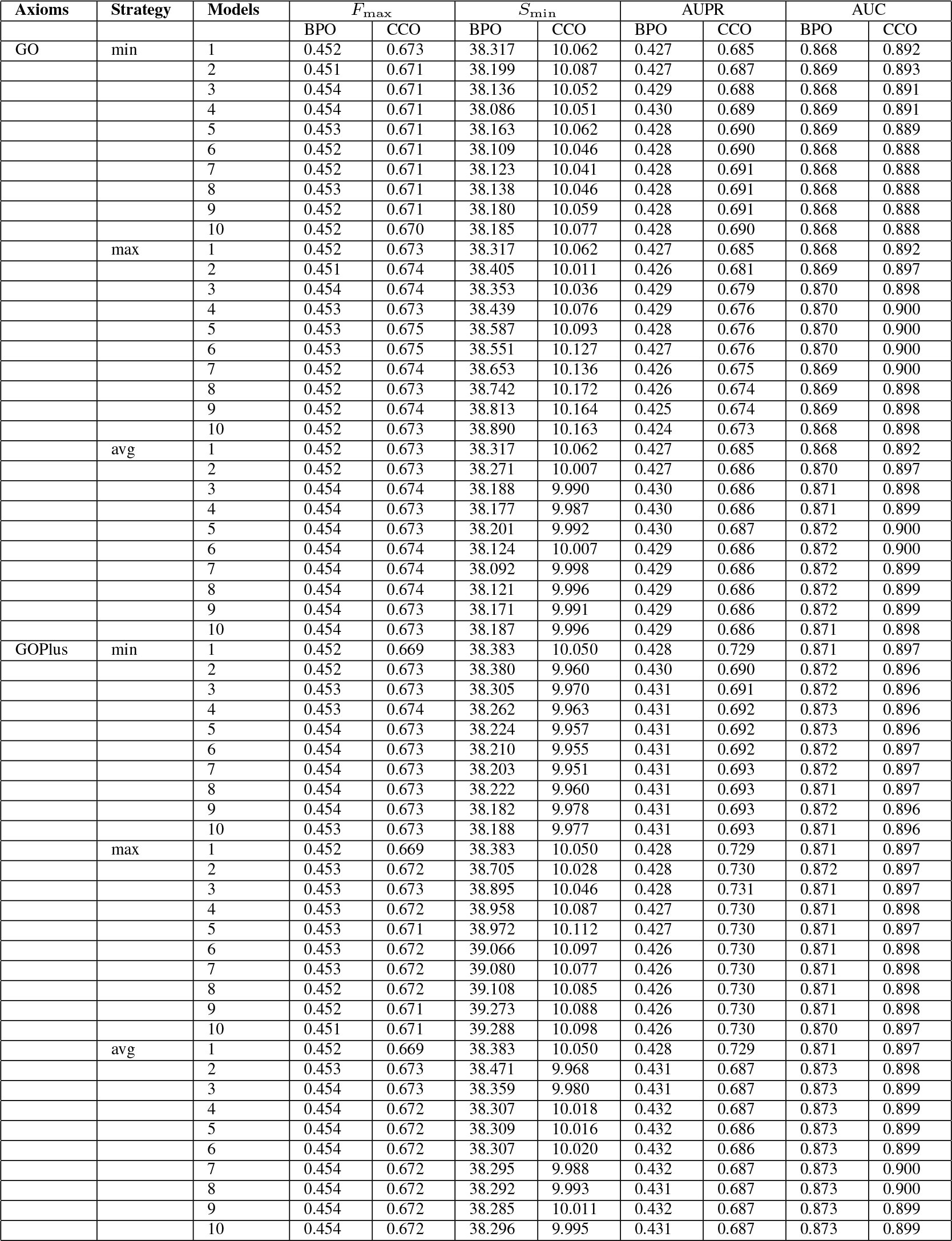
Performance of semantic entailment depending on axioms, combination strategy and number of models for DeepGOGATPredMFSE model.

**Table C5:**
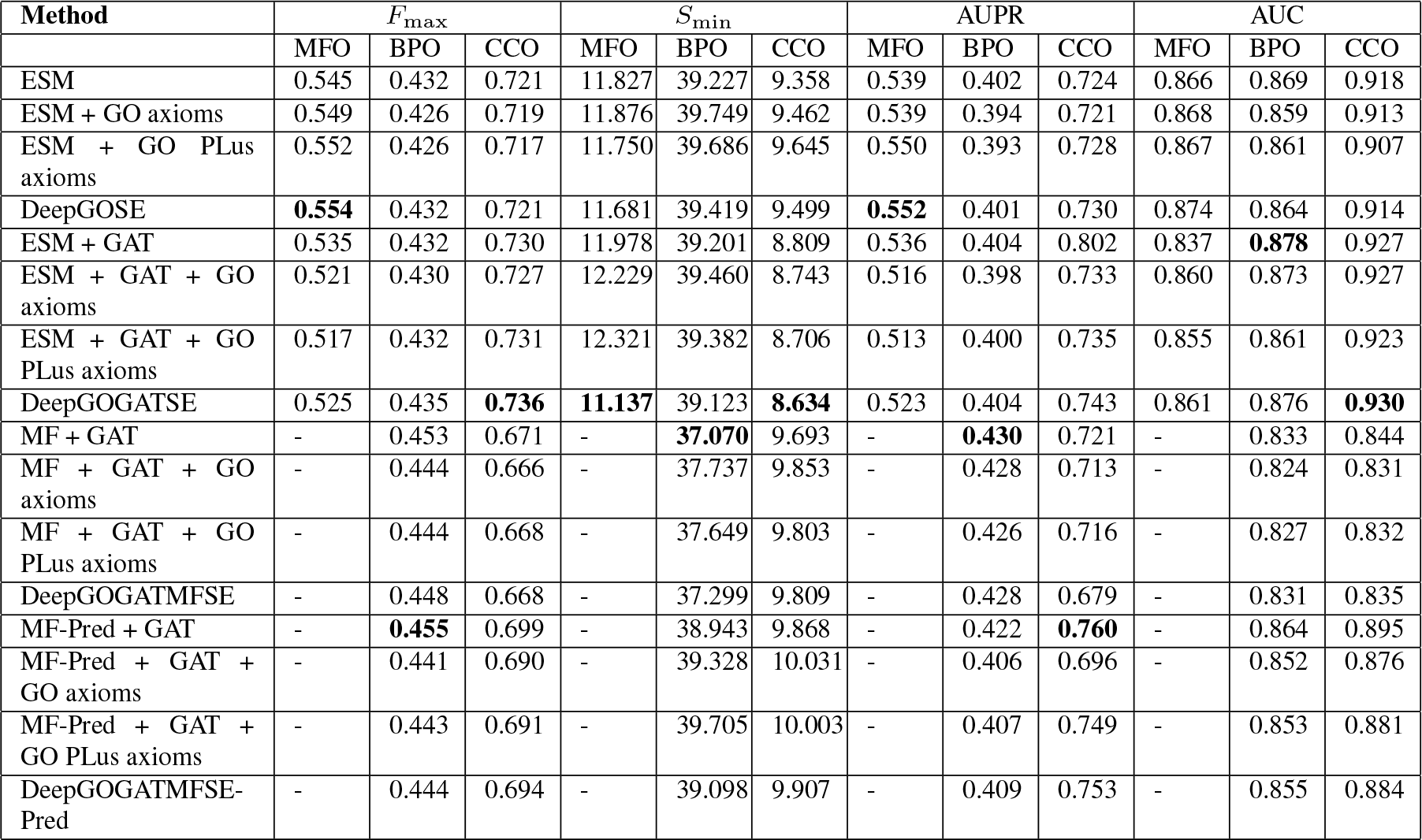
Ablation study to analyze contributions of GO and GOPlus ontology axioms, PPIs, experimental and predicted MF annotations, and Semantic Entailment to the performance.

